# Modeling Enhancer-Promoter Interactions with Attention-Based Neural Networks

**DOI:** 10.1101/219667

**Authors:** Weiguang Mao, Dennis Kostka, Maria Chikina

## Abstract

**Background:** Gene regulatory sequences play critical roles in ensuring tightly controlled RNA expression patterns that are essential in a large variety of biological processes. Specifically, enhancer sequences drive expression of their target genes, and the availability of genome-wide maps of enhancer-promoter interactions has opened up the possibility to use machine learning approaches to extract and interpret features that define these interactions in different biological contexts.

**Methods:** Inspired by machine translation models we develop an attention-based neural network model, EPIANN, to predict enhancer-promoter interactions based on DNA sequences. Codes and data are available at https://github.com/wgmao/EPIANN.

**Results:** Our approach accurately predicts enhancer-promoter interactions across six cell lines. In addition, our method generates pairwise attention scores at the sequence level, which specify how short regions in the enhancer and promoter pair-up to drive the interaction prediction. This allows us to identify over-represented transcription factors (TF) binding sites and TF-pair interactions in the context of enhancer function.

## 1. Introduction

Tightly controlled gene expression patterns are essential across a wide range of biological processes including cell differentiation, maintenance of tissue identity, and embryonic development. While the underlying mechanisms are multi-faceted and complex, cis-regulatory sequences (i.e., short, predominantly non-protein-coding DNA loci that directly affect the expression of their target genes) play critical roles. Specifically, recent research has highlighted the role of *enhancer sequences*, distal cis-regulatory elements with the capability to drive context (e.g., tissue, cell-type) specific transcription of their target genes, which are typically hundreds of kilobases away [1, 2]. Interest in these regulatory elements is also driven by the observation that DNA sequence variation in non-coding regulatory loci substantially contributes to the genetic causes of complex disease: most single nucleotide variants associated with disease are not in linkage disequilibrium with protein-coding regions, and the majority of bases in the human genome that are under negative selection are non-coding [3, 4].

Mechanistic understanding of enhancer function remains incomplete, but current models include the (context-specific) physical interaction of an enhancer with its target genes via chromatin loops, together with the binding of sequence specific transcription factors and the recruitment of coactivator proteins [1]. While uncontentious and direct experimental confirmation of enhancer function remains difficult and time-consuming, a wealth of enhancers have been annotated using comparative and functional genomics approaches coupled with bioinformatics analyses. However, without annotation of enhancers’ target genes it is difficult to infer their functional role. Fortunately genome-wide screening for enhancer-promoter interactions based on paired-end tag sequencing (ChIA-PET) or chromatin conformation capture based methods [5-7] is possible, and high-resolution Hi-C data linking regulatory sequences to promoters is publicly available for a limited number of cell-types and tissues [7].

These data open up the possibility to statistically analyze enhancer-promoter interactions (EPIs) with the goal to (i) build generalizable models that predict enhancer-promoter interaction events and to (ii) highlight informative sequences and functional genomics features in order to better understand EPI mechanisms. For instance, Whalen at al. [8] designed a method based on ensembles of boosted decision trees using functional genomics signals at enhancers, promoters and in intervening regions, and they were able to accurately predict enhancer-promoter interaction events. Subsequently, Yang et al. [9] proposed a method called PEP, demonstrating that it is possible to achieve similar predictive performance relying exclusively on sequence-based features. Here, we propose EPIANN (Enhancer-Promoter Interaction Attention-based Neural Network), which is, to the best of our knowledge, the first attention-based neural network model to predict enhancer-promoter interactions exclusively using sequence features.

Neural networks have been successfully applied in many pattern recognition tasks [10], and deep learning has become a popular tool for building DNA-sequence-based predictive models [10-14]. *Attention-based* network models were initially introduced for machine translation where they considerably improve performance[15]. More generally, the attention mechanism is broadly applicable to various matching problems, such a image captioning, and text comprehension.

Extrapolating to enhancer-promoter interaction events, given a certain enhancer segment, the attention mechanism will specify a lower level correspondence between subregions of the promoter and enhancer sequence.

In addition to predicting enhancer-promoter interactions, our model learns an attention matrix for each enhancer-promoter pair. This information can be used to identify corresponding and important sub-regions within the enhancer and promoter, respectively. Our method thereby highlights the parts of enhancer and promoter sequence that drive predictions; it allows us to analyze feature importance in the original sequence space and provides insights into the mechanism of EPI events.

## 2. Results

### 2.1. EPIANN Accurately Predicts EPI events

We compared our EPIANN method to other EPI prediction approaches: TargetFinder [8] and PEP [9]. TargetFinder uses functional genomic features such as transcription factor and histone ChIPseq, while PEP uses only sequence features, like EPIANN.

We find that our EPIANN method overall achieves comparable performance to TargetFinder and PEP, summarized in Tables 1 and 2. EPIANN outperforms the other sequence-based model, PEP, in terms of area under the receiver-operator-curve (AUROC, Table 1) though TargetFinder outperforms both sequence-only models when utilizing all functional genomics features, enhancer, promoter and in the window in-between (E/P/W). However, EPIANN performance is very similar to TargetFinder, when only enhancer and promoter features are used (E/P). In terms of area under the precision-recall-curve (AUPRC, Table 2), EPIANN performs better than TargetFinder (E/P), but not quite as good as PEP or TargetFinder (E/P/W).

**Table 1.** AUCs (Area under the ROC curve) of different enhancer-promoter interaction prediction methods for each cell line.

| Model | GM12878 | HeLa-S3 | HUVEC | IMR90 | K562 | NHEK |
| --- | --- | --- | --- | --- | --- | --- |
| EPIANN | 0.919 | 0.924 | 0.918 | 0.945 | 0.943 | 0.959 |
| PEP-Word | 0.842 | 0.843 | 0.845 | 0.898 | 0.883 | 0.917 |
| PEP-Motif | 0.866 | 0.846 | 0.839 | 0.872 | 0.892 | 0.925 |
| PEP-Integrate | 0.863 | 0.856 | 0.833 | 0.921 | 0.879 | 0.911 |
| TargetFinder (E/P) | 0.930 | 0.915 | 0.896 | 0.903 | 0.950 | 0.951 |
| TargetFinder (E/P/W) | 0.960 | 0.969 | 0.952 | 0.960 | 0.985 | 0.981 |

**Table 2.** AUPRs (Area under the precision-recall curve) of different enhancer-promoter interaction prediction methods for each cell line.

| Model | GM12878 | HeLa-S3 | HUVEC | IMR90 | K562 | NHEK |
| --- | --- | --- | --- | --- | --- | --- |
| EPIANN | 0.723 | 0.702 | 0.616 | 0.770 | 0.673 | 0.861 |
| PEP-Word | 0.807 | 0.803 | 0.760 | 0.868 | 0.836 | 0.880 |
| PEP-Motif | 0.832 | 0.820 | 0.779 | 0.732 | 0.816 | 0.892 |
| PEP-Integrate | 0.830 | 0.851 | 0.783 | 0.869 | 0.849 | 0.915 |
| TargetFinder (E/P) | 0.587 | 0.554 | 0.547 | 0.472 | 0.624 | 0.686 |
| TargetFinder (E/P/W) | 0.827 | 0.866 | 0.823 | 0.821 | 0.908 | 0.908 |

These results show that our method improves the ability to predict EPI events when only using the enhancer and promoter sequence. However, among sequence-only models PEP still outperforms EPIANN on the AUPR metric. Multiple reasons can explain this discrepancy. PEP uses considerably more information than EPIANN (PEP-Motif uses prior knowledge about TF motifs, and PEP-Word uses sequence outside the EPI loci to train its word embedding model). EPIANN also uses a smaller enhancer input regions than PEP (3 kb vs. 8 kb) which can result in missing some potentially predictive features. Moreover, EPIANN’s EPI quantification is computed as a weighted inner product between the enhancer and promoter representations in the embedding space – a relatively simple formulation. This however is by design, since we would like to force higher-order complexity to be represented in the embedding space of the promoter and enhancer sequences.

### 2.2 Decoding sequence feature importance

A key motivations for training machine learning models to predict EPIs is to give some insight into the mechanisms of these interactions. Both of the previously published models, TargetFinder and PEP, provided some mechanistic insight by using feature importance analysis. However, both TargetFinder and PEP use only feature occurrence profiles as their input, discarding spatial information. In contrast, we designed EPIANN with location-based feature decoding in mind. It directly reports an attention-based importance score for each position combination between the enhancer and promoter (i.e., an attention matrix) of an analyzed EPI event. This information can be used to compile importance scores, delineate important sub-regions, and highlight meaningful sequence annotation features.

#### Attention regions highlight sequence annotation features

In order to analyze feature importance at the sequence level we label each base in the enhancer and promoter region with its marginal attention (row-wise or column-wise maximum of the full attention matrix) to create an attention track. The attention track can be visualized in a genome browser giving a detailed view of individual promoter and enhancer features. We found that the attention regions generally correlate well with other genome annotations such as transcription factor motifs and DNAseq footprinting signal, which is a measure of TF occupancy (see Methods for footprinting details). An illustrative example is shown in Figure 1.

**Figure 1.**
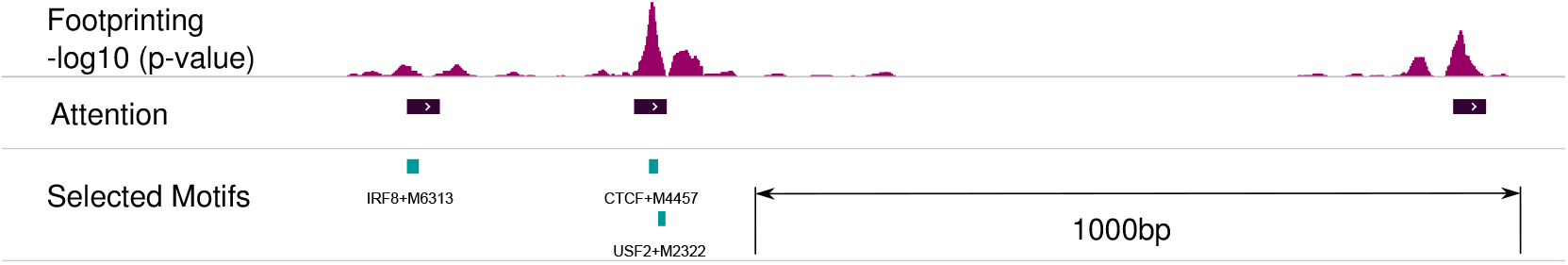
Attention regions can be used to generate a feature importance track that can be inspected visually and aligned with other genomic information.

#### Attention regions highlight transcription factors contributing to EPI events

Attention regions can also be used for statistical feature importance analysis. We can quantify the over-representation of TF motifs within top scoring attention regions relative to the entire input sequence. For this analysis, we used the top 5 regions of length 45bp (that corresponds to the sum of filter width (15) and the maxpool size (30)). Results for the IMR90 cell line are depicted in Figure 2. Consistent with results reported based on feature importance analysis form PEP and TargetFinder we observe a role for CTCF and EGR family members. However, we also detect new signals. For example, one of our most consistent findings is that motifs for the NRF1 transcription factor are strongly over-represented in enhancer sequences. NRF1, also known as nuclear respiratory factor-1, is a broadly expressed TF that activates genes involved in respiration and mitochondrial biogenesis. IMR90 is a contractile muscle-like cell line and NRF1 is particularly important for muscle biology, due to muscle cells’ unique energy requirements [16]. Neither TargetFinder nor PEP makes a similar prediction. Whether or not NRF1 mediates EPIs must be determined with experimental follow up, however the observation highlights an important feature of our method. Since EPIANN feature importance is based on explicitly specified attention region coordinates, EPI loci can be analyzed directly. Therefore, any and all TF motifs (or any other sequence based analysis) can be used *after* training the model to assess feature importance.

**Figure 2.**
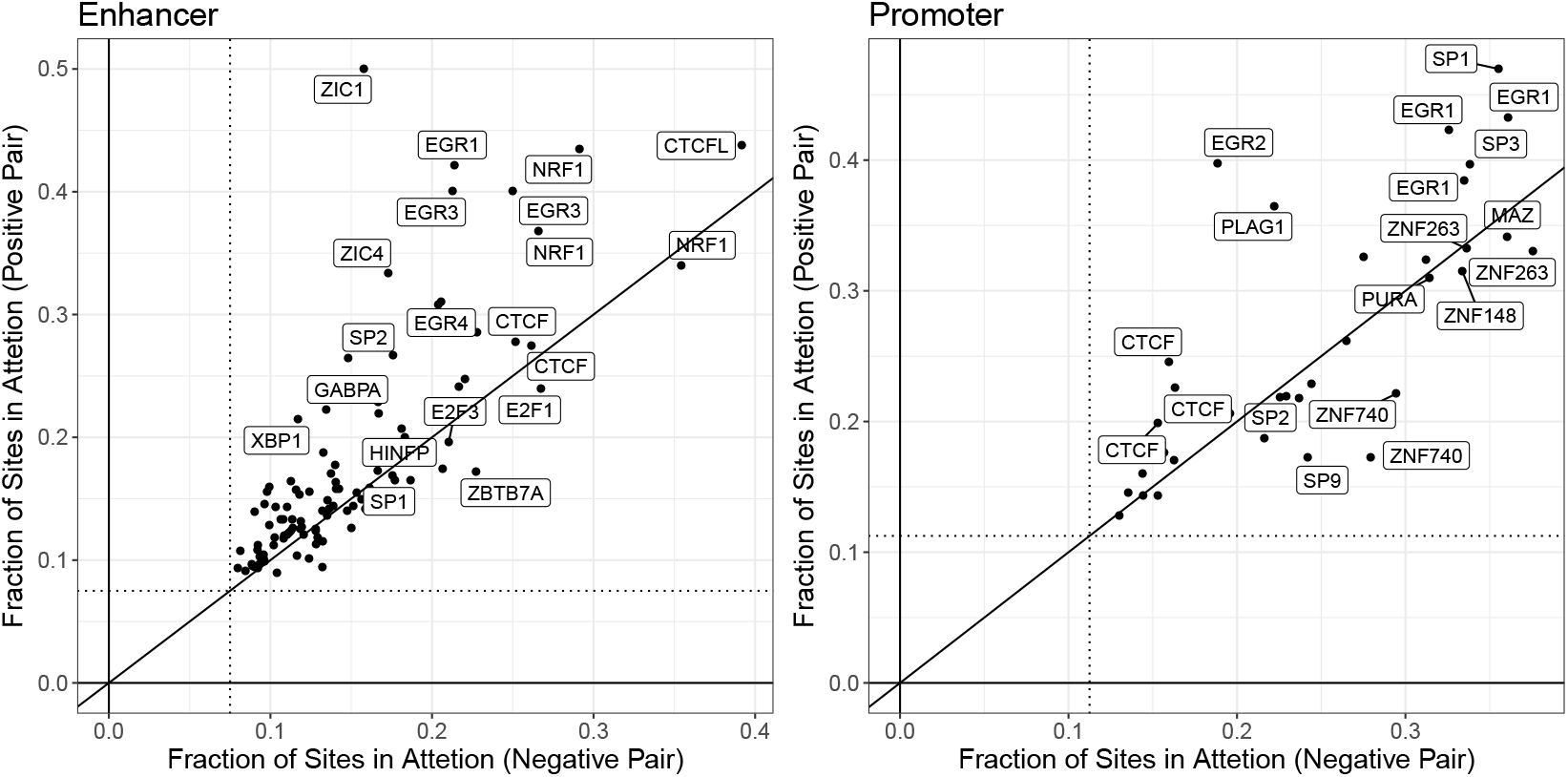
Enrichment of transcription factor motif sites within the attention regions. Enrichment is quantified as the fraction of all possible sites in the input enhancer or promoter regions that are captured in the smaller attention region. The dotted line specifies the expected fraction for randomly selected mock “attention” regions. Comparing the degree of enrichment in attention regions learned for positive and negative interaction examples we find that the enrichments show similar signals (but for different TFs), though especially in the case of enhancers some TFs are more enriched when only positive examples are considered. Transcription factors may appear more than once due to multiple available motifs.

In the case of TargetFinder, the feature importance is limited by the input data types and no NRF1 ChIPseq was available. PEP uses known motifs, which we presume included at least one NRF1, but their feature analysis did not deem it important. This difference may arise from the exact motif used, as several ones are available.

#### Attention regions correlate with transcription factor occupancy

Using attention regions visualized as genome tracks we noticed a striking correspondence between DNAase footprinting, which measures transcription factor occupancy, and out attention scores. Summarizing the trend genome-wide we find that attention regions are highly biased towards occupied sites (see Figure 3A). Moreover, intersecting motif positions with occupancy status we find that some motifs show much stronger enrichment in attention regions specifically for their occupied sites (though this observation is specific for enhancer sequences). This demonstrates that attention scores provide information beyond simple motif matches, which are captured in the first convolution layers of the neural network, but reflects information from the entire model which can specify higher-order interactions that correlate with whether or not a TF consensus site is actually occupied.

**Figure 3.**
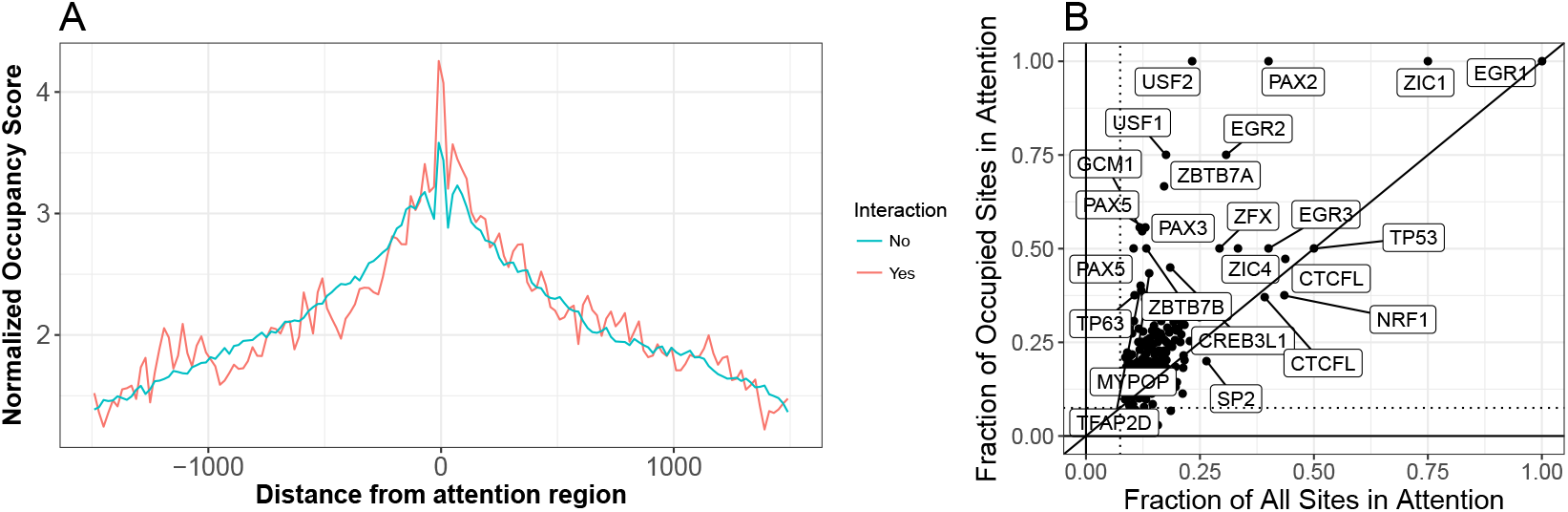
(A) Distribution of normalized occupancy scores is centered on top attention regions. The raw occupancy scores are –log_10_ p-values reported by pyDNAse[17, 18]. In order to compute the normalized profile we only consider input regions that have a minimum occupancy score of 3, and among those the profile is normalized so that the maximum is equal to the median maximum among all regions. This normalization is important since the p-value depends directly on the number of reads in the region and the normalization ensures that we do not compare the read depth but only the overall association of attention regions with occupancy status. (B) Comparing fraction of all sites captured in the attention with the fraction of occupied sites within enhancer regions. Some factors are more strongly enriched if only occupied sites are considered, demonstrating that attention mechanism provides feature importance assessment that goes beyond simple motif matching.

#### Attention regions suggest transcription factor interactions

So far our analysis has only considered individual promoter and enhancer sequences. However, since the attention regions correspond to a specific sequence instances matched within each enhancer-promoter *pair*, we can also ask if there are pairs of transcription factors whose motifs contribute together (one in the enhancer sequence and one in the promoter sequence) to a positive EPI prediction. We do this by comparing the distribution of TF-pairs in the top attention regions of interacting and non-interacting promoter-enhancer pairs. The TF-pairs enriched in the attention of interacting promoter-enhancer pairs are depicted in an enrichment heatmap (Figure 4).

**Figure 4.**
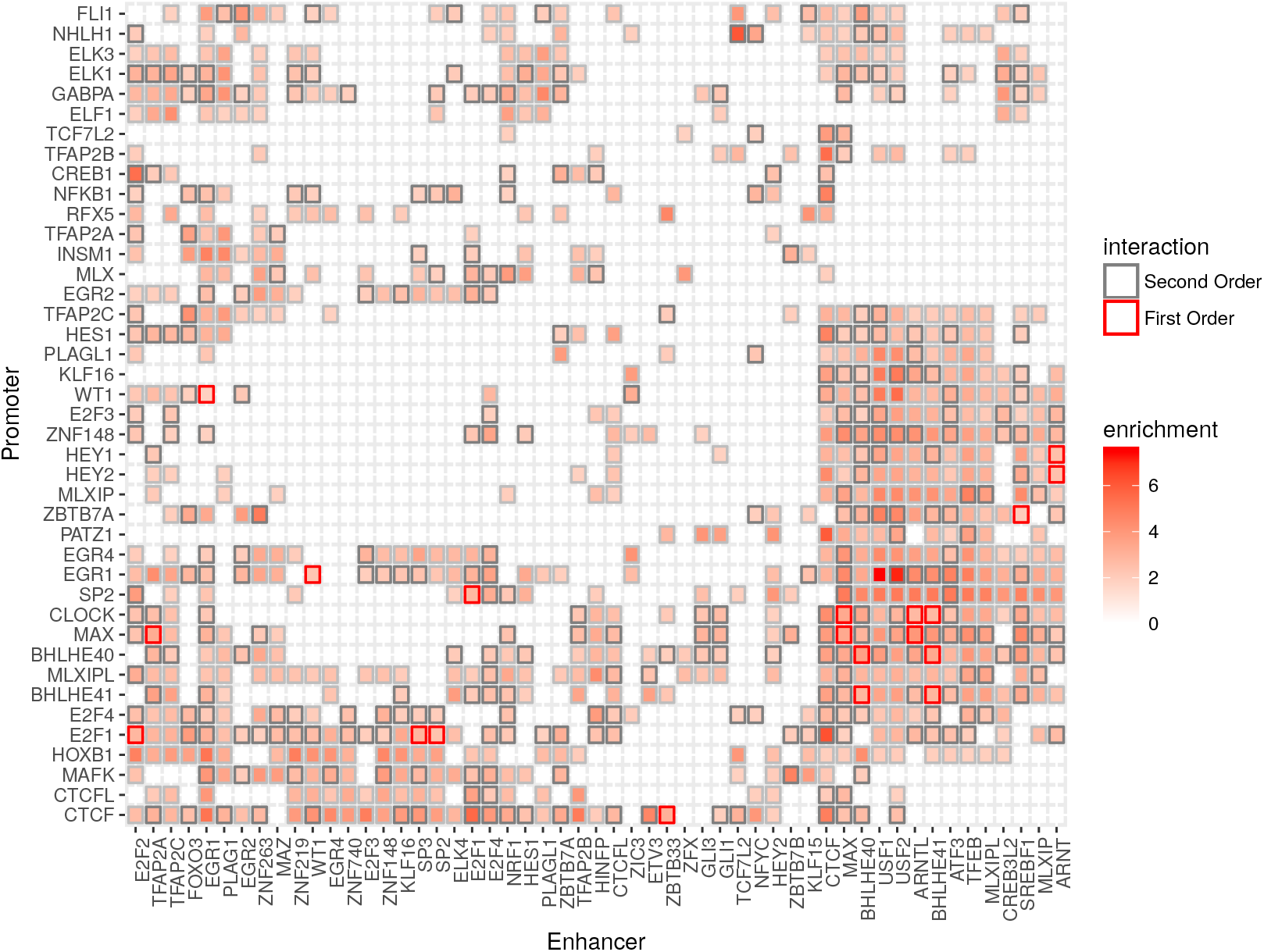
Enrichment of transcription factor motif within the attention regions of interacting enhancer-promoter pairs. Since multiple motifs are available for each transcription factor we take the maximum enrichment among all possible motif combinations. Border color indicate the existence of known first-order (direct) or second-order interactions in BioGRID [19]

We find that most interactions are heterogeneous, involving two different TFs. The enrichment map is also highly asymmetric; that is, for many enriched TF-pairs the participating factors have motifs in only the enhancer or the promoter sequencing of EPI loci. For example even though EGR2 and CTCF are some of the strongest single factor enrichments in both promoters and enhancers, the interaction is only enriched when EGR2 is in the enhancer and CTCF is in the promoter (see Figure 4). We also note a trend of multiple clusters of varying sizes where almost all pair-wise interactions appear enriched.

We speculate that these TF-clusters participate either in direct or higher order interactions that drive the formation of an enhancer-promoter interaction complex. We find that the enrichment score is indeed predictive of both known first and known second-order protein-protein interactions from BioGRID [19]. We find that the 50 most enriched TF pairs have a 0.03 probability of physically interacting compared with a baseline of 0.01 among all tested pairs. The corresponding numbers for second-order interactions are 0.49 (50 most enriched) and 0.29 (baseline). The enrichment of known interactions suggests that our model learns a meaningful biological representation and can be used to form hypotheses about new interactions that mediate EPI events–for example, our top scoring interaction is between USF1/USF2 and EGR1. While we know of no data supporting a direct interaction between these genes, the USF1-EGR1 interaction has been previously suggested based on overlapping patterns of ChIPseq signals [20].

## 3. Discussion

The mechanism of enhancer-promoter interactions is of tremendous interest but is currently poorly understood. Even though it is clear that EPI events can be predicted from sequence features, it is not yet possible to use this to predict new EPIs as all the predictions are highly cell-type specific. That is, it is not yet possible to accurately predict EPIs for a tissue/cell-line that is different from the one used to train the model. Thus, the models do not replace the need for experimental HiC data. However, there is hope that the models can provide mechanistic insight into the nature of the EPI complex via feature importance analysis. Our model is designed with an attention mechanism, which provides single and pairwise importance scores for each position in the enhancer and promoter input regions making it feasible to analyze feature importance in more detail.

## 4 Methods

### 4.1. Data Preparation

We utilize the same EPI data originally collected by TargetFinder, so we can make a direct comparison with TargetFinder and PEP. The EPI data contains six cell lines (GM12878, HeLa-S3, HUVEC, IMR90, K562 and NHEK), for each cell line it records the genomic coordinates of enhancers and promoters with indicators of EPIs. The ratio of positive interaction pairs to negative interaction pairs is about one to twenty, which is common for Hi-C data. To overcome this imbalanced training problem, we augment the positive data to twenty folds. The preprocessing pipeline for each cell line contains the following steps:

1. Start from the imbalanced data *D*.
2. Split D into a training set *D_train_* (90% of *D*) and a test set *D_test_* (10% of *D*) by stratified sampling.
3. Augment *D_train_* to get a balanced dataset *D_aug_*.
4. Train the model on *D_aug_*.
5. Evaluate the model on *D_test_*.

The inputs for the model are two extended DNA segments which contains the annotated enhancers and promoters correspondingly. The length of the extended window is chosen as 3K bp for enhancer and 2K bp for promoter, which try to capture all the relevant regions around enhancers and promoters. During the augmentation, we slide the extended region around the enhancer or promoter as long as it contains most of the functional parts. We fix *D_test_* and *D_train_* for each cell line in order to compare the performance with TargetFinder and PEP. The results are reported in Tables 1 and 2.

### 4.2. Attention-Based Neural Network Model Architecture

We propose a neural network structure to predict enhancer-promoter interactions only using sequence-based features. The overall network structure is shown in Figure 5 and there are three functional blocks of the models which are **attention mechanism**, **interaction quantification** and **multi-task learning**.

**Figure 5.**
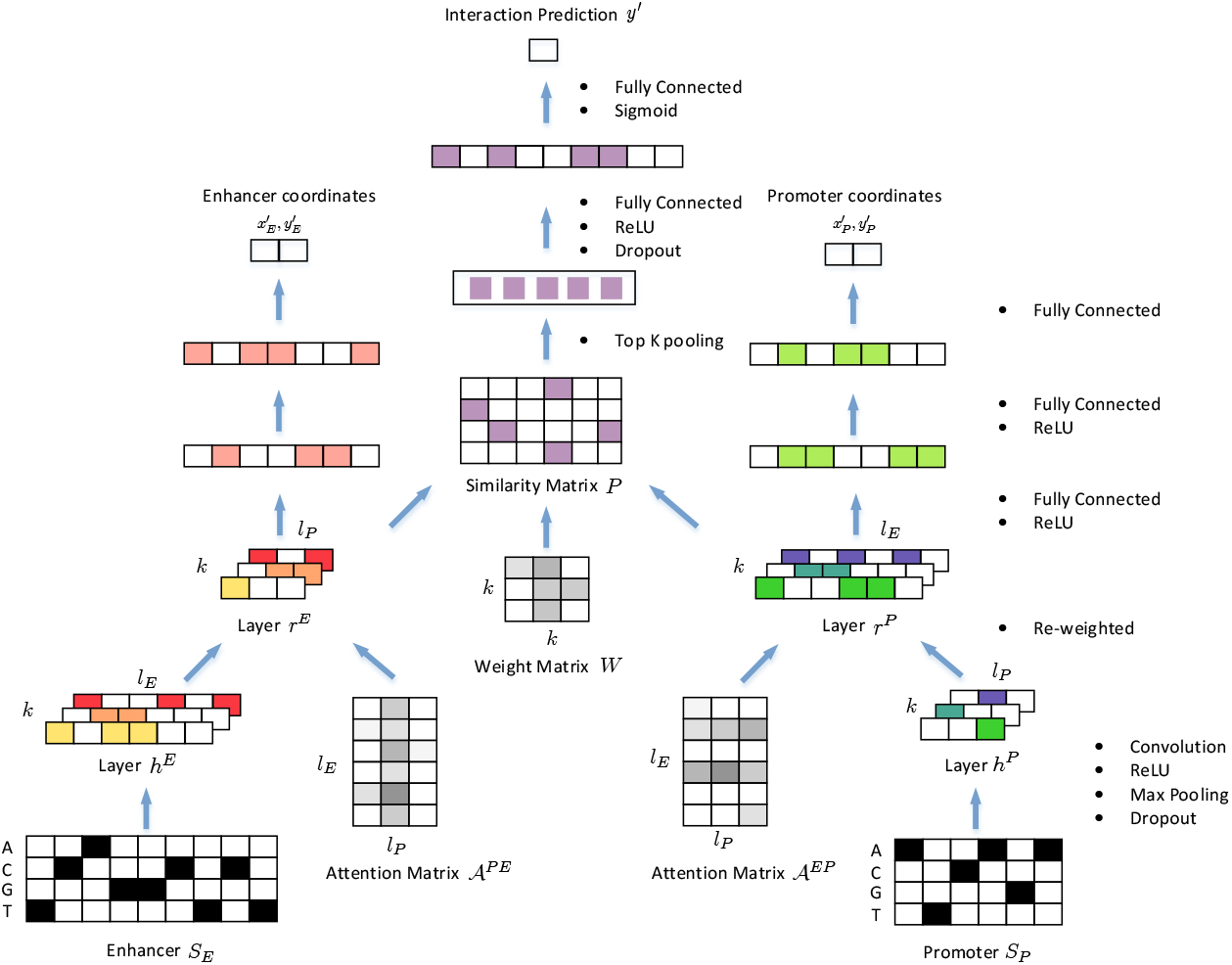
Schematic overview of EPI. For visualization, the parameters are shrunk accordingly. The length of extended enhancer *S_E_* is set to be 9 and the length of extended promoter *S_P_* is set to be 6. After passing *S_E_* and *S_P_* through the convolution layer with *k* = 3 kernels and the max pooling layer, the corresponding dimension of *h^E^* becomes ℝ*^l_E_ × k^*, where *l_E_* = 6. Similarly the dimension of *h^p^* becomes ℝ*^l_P_ × k^*, where *l_P_* = 3.

#### 4.2.1. Attention Mechanism

The two extended DNA segments will be transformed into one-hot encoding with four channels (A, C, G and T). After embedding, enhancer and promoter sequences *S_E_* and *S_P_* are passed through separate convolutional layers which share the same *k* filters with outputs denoted as *h^E^* ∈ ℝ*^l_E_ × k^* and *h^P^ ∈ ℝ^l_p_ × k^*. Convolutional kernels are equivalent as position specific scoring matrice[21], by which local sequence patterns are encoded in *h^E^* and *h^P^*. The next layers *r^E^* and *r^P^* are computed as weighted sums of *h_E_* and *h_P_* correspondingly.

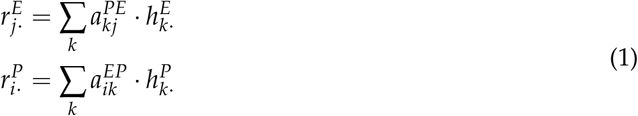

The weight 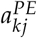 and 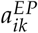 are computed by

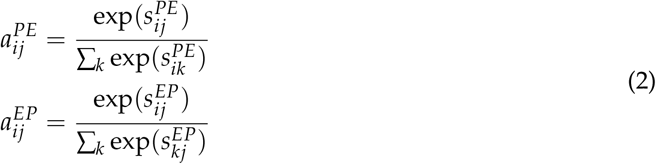

where

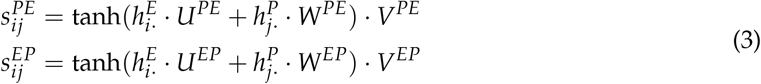

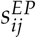 shows how well *h^E^* around position *i* align with *h_p_* at position *j*. 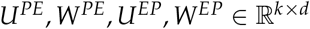 and 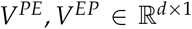 are all hidden variables, and *d* represents the hidden dimension which is a hyperparameter in the model. The probability 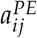 reflects the importance of 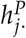 with respect to all possible 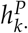 given 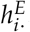. Similarly the probability 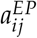 represents the importance of 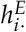 regarding all possible 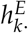 given 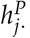. This formulation of alignment is called soft attention[15, 22]. We denote these two weight/attention matrices as 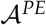 and 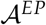, and scoring matrices as 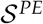 and 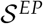. They all come with the same dimension ℝ*^l_E_ × l_P_^*.

#### 4.2.2 Interaction Quantification

After enhancers and promoters are projected into the same embedding space as *r^E^* and *r^p^*, we would like to calculate the weighted inner product between corresponding embeddings, which is a similarity measure of embedding vectors. In this way, we interpret the probability of interaction event at sequence level to be the similarity level of embeddings after projection. If the paired embedding are aligned really well, it means these pairs of interactions can lead to the interaction at sequence level. Similar ideas have been use to model the interactions between semantic segment pairs[23], correspondences between images and captions [24, 25], etc. *W ∈ ℝ^k × k^* represents the weight matrix which is a free parameter in the model. We define the similarity matrix as *P ∈ ℝ^l_P_ × l_E_^*.

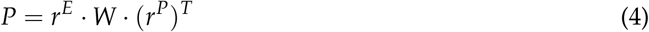

Then we pass *P* through top *K*-pooling layer, a fully connected layer with ReLU activation and a fully connected layer with Sigmoid activation. The final output is *y*′ which is a probability indicating the chance that two sequences will interact. We define binary cross-entropy loss on *y*′.

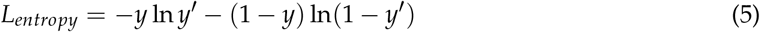

where *y* represents the true label which is 0 or 1.

#### 4.2.3. Multi-task Learning

Multi-task learning idea has been incorporated into a large number of deep learning frameworks [26]. Multi-task learning can force the model to learn more generalizable features regarding multiple related tasks at the same time and it can be regarded as an implicit regularization on the model. The input sequences *S_E_* and *S_P_* to this model are not exactly the enhancer and promoter sequences but a larger windows which contains the enhancer and promoter regions. Other than the interaction prediction task, we can also introduce one additional task which is to infer the enhancer and promoter regions from the extended segments. We call this task *coordinate prediction*. This idea is mainly inspired by the objection detection work in the computer vision field. R-CNN[27], Fast R-CNN [28] and Faster R-CNN [29] are proposed to localize and label the boundaries for detected object in the images.

We formulate *coordinate prediction* task by asking the model to predict internal coordinates of functional regions for *S_E_* and *S_P_*. After passing *r^E^* and *r^P^* through separate fully connected layers with ReLU activations, we will get regression results of internal enhancer coordinates 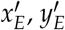 and internal promoter coordinates 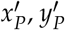. We define *l_2_* loss *L_coor_* on 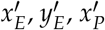, and 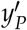.

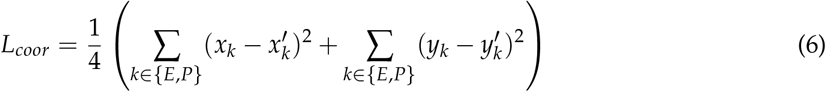

where *x_E_*, *y_E_*, *x_P_* and *y_P_* are the true internal coordinates we generate when augmenting the data. Thus the overall loss function is defined as

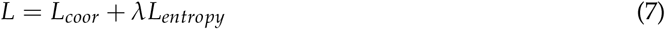

where *λ* is a hyperparameter. All hyperparameters are tuned by cross validation on *D_train_*. For training, we used adam optimizer [30] as the stochastic gradient descent optimizer with a learning rate of 1*e* – 3 and it takes 90 epochs. Early stopping [31] and dropout [32] are incorporated into the neural network structure and the optimization process.

### 4.3. Motif analysis

We scanned the genome for putative TF binding sites using position weight matricies from Transfac [33] and the genome scanning tool provided by Homer [34].

### 4.4. DNAseq footprintg

Footprinting is a standard analysis of DNase signal that goes beyond calling open regions to find smaller regions that are depleted for DNase cuts due to their occlusion by a DNA binding factor. We used the pyDNAse implementation [17, 18], to quantify the extent to which specific sites in the genome are occupied by DNA binding proteins.

## Acknowledgments

NSF 1458766, NIH BD2K U54HG008540, NIH HG00854003, NIH MH109009, D.K. was supported by the National Institutes of Health [1R01GM115836]

## Abbreviations

The following abbreviations are used in this manuscript:

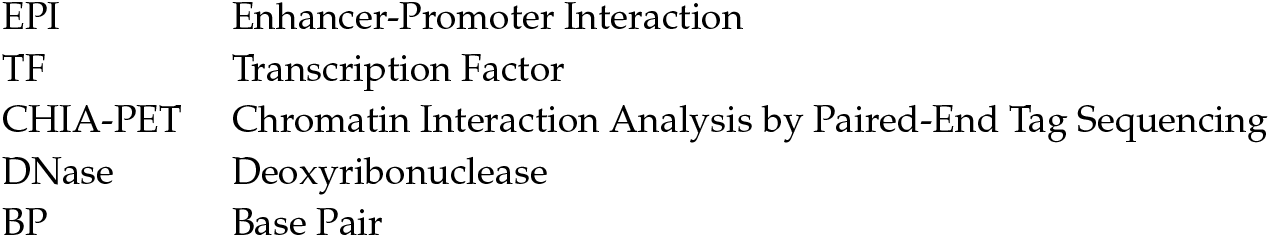

## Appendix A *F*_1_ **scores**

**Table A1.** *F*_1_ scores of different enhancer-promoter interaction prediction methods for each cell line.

| MODEL | GM12878 | HeLa-S3 | HUVEC | IMR90 | K562 | NHEK |
| --- | --- | --- | --- | --- | --- | --- |
| EPIANN | 0.699 | 0.639 | 0.590 | 0.711 | 0.626 | 0.797 |
| PEP-WORD | 0.784 | 0.782 | 0.749 | 0.854 | 0.826 | 0.874 |
| PEP-MOTIF | 0.816 | 0.796 | 0.762 | 0.725 | 0.810 | 0.887 |
| PEP-INTEGRATE | 0.813 | 0.824 | 0.761 | 0.865 | 0.836 | 0.902 |
| TARGETFINDER (E/P) | 0.489 | 0.496 | 0.448 | 0.391 | 0.486 | 0.675 |
| TARGETFINDER (E/P/W) | 0.773 | 0.787 | 0.748 | 0.683 | 0.795 | 0.872 |

## Appendix B Cross-Validation Performance

A more comprehensive way to compare different models it to report the cross-validation performance. But it is not feasible to do that with any neural network model due to the computational cost. Thus we fix *D_train_* and *D_test_* and only report performance based on these two randomly selected sets for each cell line. In order to show the reported performance is not biased regarding the choice of *D_train_* and *D_test_*, we list the cross-validation performance of all submodels of PEP and TargetFinder.

**Table A2.** The mean *F_1_* scores of different enhancer-promoter interaction prediction methods for each cell line regarding 10-fold cross validation. The values in the parentheses are the corresponding standard deviations.

| Model | GM12878 | HeLa-S3 | HUVEC | IMR90 | K562 | NHEK |
| --- | --- | --- | --- | --- | --- | --- |
| PEP-Motif | 0.846 | 0.820 | 0.728 | 0.809 | 0.797 | 0.837 |
| PEP-Word | 0.837 | 0.809 | 0.722 | 0.820 | 0.780 | 0.834 |
| PEP-Integrate | 0.851 | 0.821 | 0.743 | 0.834 | 0.795 | 0.847 |
| TargetFinder (E/P) | 0.516(0.048) | 0.541(0.023) | 0.405(0.027) | 0.413(0.032) | 0.536(0.042) | 0.540(0.034) |
| TargetFinder (EE/P) | 0.835(0.014) | 0.817(0.015) | 0.671(0.017) | 0.808(0.029) | 0.798(0.019) | 0.819(0.016) |
| TargetFinder (E/P/W) | 0.763(0.033) | 0.853(0.022) | 0.712(0.024) | 0.731(0.039) | 0.821(0.024) | 0.864(0.022) |

**Table A3.** The mean AUCs (Area under the ROC curve) of different enhancer-promoter interaction prediction methods for each cell line regarding 10-fold cross validation. The values in the parentheses are the corresponding standard deviations.

| Model | GM12878 | HeLa-S3 | HUVEC | IMR90 | K562 | NHEK |
| --- | --- | --- | --- | --- | --- | --- |
| PEP-Motif | 0.963 | 0.967 | 0.941 | 0.951 | 0.956 | 0.966 |
| PEP-Word | 0.966 | 0.966 | 0.945 | 0.958 | 0.958 | 0.973 |
| PEP-Integrate | 0.967 | 0.972 | 0.947 | 0.961 | 0.960 | 0.975 |
| TargetFinder (E/P) | 0.926 (0.014) | 0.935(0.012) | 0.896(0.005) | 0.906(0.015) | 0.929(0.012) | 0.941(0.008) |
| TargetFinder (EE/P) | 0.971(0.008) | 0.967(0.008) | 0.939(0.007) | 0.966(0.013) | 0.965(0.007) | 0.971(0.010) |
| TargetFinder (E/P/W) | 0.964(0.010) | 0.975(0.009) | 0.957(0.008) | 0.964 (0.010) | 0.964(0.013) | 0.983(0.009) |

**Table A4.**
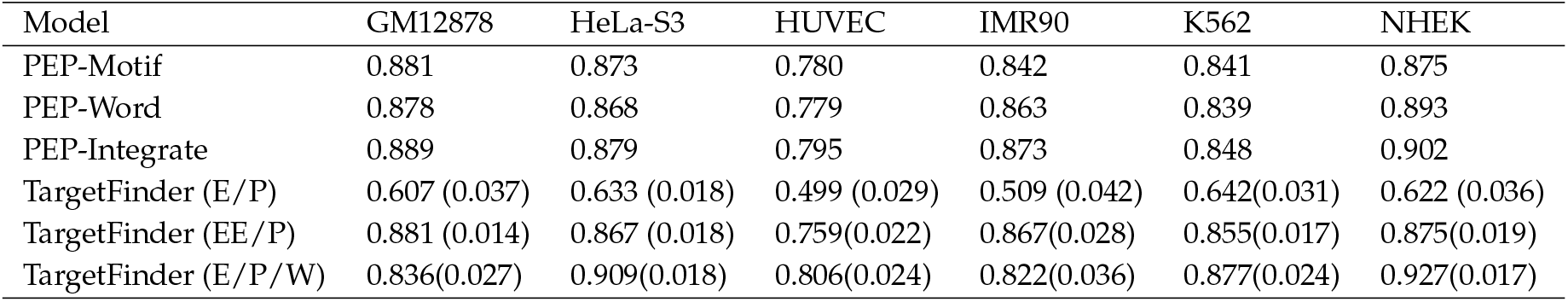
The mean AUPRs (Area under the precision-recall curve) of different enhancer-promoter interaction prediction methods for each cell line regarding 10-fold cross validation. The values in the parentheses are the corresponding standard deviations.

| Model | GM12878 | HeLa-S3 | HUVEC | IMR90 | K562 | NHEK |
| --- | --- | --- | --- | --- | --- | --- |
| PEP-Motif | 0.881 | 0.873 | 0.780 | 0.842 | 0.841 | 0.875 |
| PEP-Word | 0.878 | 0.868 | 0.779 | 0.863 | 0.839 | 0.893 |
| PEP-Integrate | 0.889 | 0.879 | 0.795 | 0.873 | 0.848 | 0.902 |
| TargetFinder (E/P) | 0.607 (0.037) | 0.633 (0.018) | 0.499 (0.029) | 0.509 (0.042) | 0.642(0.031) | 0.622 (0.036) |
| TargetFinder (EE/P) | 0.881 (0.014) | 0.867 (0.018) | 0.759(0.022) | 0.867(0.028) | 0.855(0.017) | 0.875(0.019) |
| TargetFinder (E/P/W) | 0.836(0.027) | 0.909(0.018) | 0.806(0.024) | 0.822(0.036) | 0.877(0.024) | 0.927(0.017) |

## References

1. Long, H.K.; Prescott, S.L.; Wysocka, J. Ever-changing landscapes: transcriptional enhancers in development and evolution. Cell 2016, 167, 1170–1187.

2. Spitz, F. Gene regulation at a distance: from remote enhancers to 3D regulatory ensembles. Seminars in cell & developmental biology. Elsevier, 2016, Vol. 57, pp. 57–67.

3. Albert, F.W.; Kruglyak, L. The role of regulatory variation in complex traits and disease. Nature Reviews Genetics 2015, 16, 197–212.

4. Huang, Y.F.; Gulko, B.; Siepel, A. Fast, scalable prediction of deleterious noncoding variants from functional and population genomic data. Nature genetics 2017, 49, 618–624.

5. Fullwood, M.J.; Liu, M.H.; Pan, Y.F.; Liu, J.; Xu, H.; Mohamed, Y.B.; Orlov, Y.L.; Velkov, S.; Ho, A.; Mei, P.H.; others. An oestrogen-receptor-α-bound human chromatin interactome. Nature 2009, 462, 58–64.

6. Dostie, J.; Richmond, T.A.; Arnaout, R.A.; Selzer, R.R.; Lee, W.L.; Honan, T.A.; Rubio, E.D.; Krumm, A.; Lamb, J.; Nusbaum, C.; others. Chromosome Conformation Capture Carbon Copy (5C): a massively parallel solution for mapping interactions between genomic elements. Genome research 2006, 16, 1299–1309.

7. Rao, S.S.; Huntley, M.H.; Durand, N.C.; Stamenova, E.K.; Bochkov, I.D.; Robinson, J.T.; Sanborn, A.L.; Machol, I.; Omer, A.D.; Lander, E.S.; others. A 3D map of the human genome at kilobase resolution reveals principles of chromatin looping. Cell 2014, 159, 1665–1680.

8. Whalen, S.; Truty, R.M.; Pollard, K.S. Enhancer-promoter interactions are encoded by complex genomic signatures on looping chromatin. Nature genetics 2016, 48, 488.

9. Yang, Y.; Zhang, R.; Singh, S.; Ma, J. Exploiting sequence-based features for predicting enhancer-promoter interactions 2017.

10. Min, S.; Lee, B.; Yoon, S. Deep learning in bioinformatics. Briefings in bioinformatics 2016, p. bbw068.

11. Ching, T.; Himmelstein, D.S.; Beaulieu-Jones, B.K.; Kalinin, A.A.; Do, B.T.; Way, G.P.; Ferrero, E.; Agapow, P.M.; Xie, W.; Rosen, G.L.; others. Opportunities And Obstacles For Deep Learning In Biology And Medicine. bioRxiv 2017, p. 142760.

12. Alipanahi, B.; Delong, A.; Weirauch, M.T.; Frey, B.J. Predicting the sequence specificities of DNA-and RNA-binding proteins by deep learning. Nature biotechnology 2015, 33, 831–838.

13. Zhou, J.; Troyanskaya, O.G. Predicting effects of noncoding variants with deep learning–based sequence model. Nature methods 2015, 12, 931.

14. Poplin, R.; Newburger, D.; Dijamco, J.; Nguyen, N.; Loy, D.; Gross, S.S.; McLean, C.Y.; DePristo, M.A. Creating a universal SNP and small indel variant caller with deep neural networks. bioRxiv 2016, p. 092890.

15. Bahdanau, D.; Cho, K.; Bengio, Y. Neural machine translation by jointly learning to align and translate. arXiv preprint arXiv:1409.0473 2014.

16. Ramachandran, B.; Yu, G.; Gulick, T. Nuclear respiratory factor 1 controls myocyte enhancer factor 2A transcription to provide a mechanism for coordinate expression of respiratory chain subunits. Journal of Biological Chemistry 2008, 283, 11935–11946.

17. Piper, J.; Elze, M.C.; Cauchy, P.; Cockerill, P.N.; Bonifer, C.; Ott, S. Wellington: a novel method for the accurate identification of digital genomic footprints from DNase-seq data. Nucleic acids research 2013, 41, e201–e201.

18. Piper, J.; Assi, S.A.; Cauchy, P.; Ladroue, C.; Cockerill, P.N.; Bonifer, C.; Ott, S. Wellington-bootstrap: Differential DNase-seq footprinting identifies cell-type determining transcription factors. BMC genomics 2015, 16, 1000.

19. Chatr-aryamontri, A.; Oughtred, R.; Boucher, L.; Rust, J.; Chang, C.; Kolas, N.K.; O’Donnell, L.; Oster, S.; Theesfeld, C.; Sellam, A.; others. The BioGRID interaction database: 2017 update. Nucleic acids research 2017, 45, D369–D379.

20. Giannopoulou, E.G.; Elemento, O. Inferring chromatin-bound protein complexes from genome-wide binding assays. Genome research 2013, 23, 1295–1306.

21. Stormo, G.D.; Schneider, T.D.; Gold, L.; Ehrenfeucht, A. Use of the ‘Perceptron’algorithm to distinguish translational initiation sites in E. coli. Nucleic acids research 1982, 10, 2997–3011.

22. Xu, K.; Ba, J.; Kiros, R.; Cho, K.; Courville, A.; Salakhudinov, R.; Zemel, R.; Bengio, Y. Show, attend and tell: Neural image caption generation with visual attention. International Conference on Machine Learning, 2015, pp. 2048–2057.

23. Liu, P.; Qiu, X.; Huang, X. Modelling Interaction of Sentence Pair with coupled-LSTMs. arXiv preprint arXiv:1605.05573 2016.

24. Karpathy, A.; Joulin, A.; Li, F.F.F. Deep fragment embeddings for bidirectional image sentence mapping. Advances in neural information processing systems, 2014, pp. 1889–1897.

25. Karpathy, A.; Fei-Fei, L. Deep visual-semantic alignments for generating image descriptions. Proceedings of the IEEE Conference on Computer Vision and Pattern Recognition, 2015, pp. 3128–3137.

26. Ruder, S. An Overview of Multi-Task Learning in Deep Neural Networks. arXiv preprint arXiv:1706.05098 2017.

27. Girshick, R.; Donahue, J.; Darrell, T.; Malik, J. Rich feature hierarchies for accurate object detection and semantic segmentation. Proceedings of the IEEE conference on computer vision and pattern recognition, 2014, pp. 580–587.

28. Girshick, R. Fast r-cnn. Proceedings of the IEEE international conference on computer vision, 2015, pp. 1440–1448.

29. Ren, S.; He, K.; Girshick, R.; Sun, J. Faster R-CNN: Towards real-time object detection with region proposal networks. Advances in neural information processing systems, 2015, pp. 91–99.

30. Kingma, D.; Ba, J. Adam: A method for stochastic optimization. arXiv preprint arXiv:1412.6980 2014.

31. Prechelt, L. Early stopping-but when? Neural Networks: Tricks of the trade 1998, pp. 553–553.

32. Srivastava, N.; Hinton, G.E.; Krizhevsky, A.; Sutskever, I.; Salakhutdinov, R. Dropout: a simple way to prevent neural networks from overfitting. Journal of Machine Learning Research 2014, 15, 1929–1958.

33. Matys, V.; Kel-Margoulis, O.V.; Fricke, E.; Liebich, I.; Land, S.; Barre-Dirrie, A.; Reuter, I.; Chekmenev, D.; Krull, M.; Hornischer, K.; others. TRANSFAC^®^ and its module TRANSCompel^®^: transcriptional gene regulation in eukaryotes. Nucleic acids research 2006, 34, D108–D110.

34. Heinz, S.; Benner, C.; Spann, N.; Bertolino, E.; Lin, Y.C.; Laslo, P.; Cheng, J.X.; Murre, C.; Singh, H.; Glass, C.K. Simple combinations of lineage-determining transcription factors prime cis-regulatory elements required for macrophage and B cell identities. Molecular cell 2010, 38, 576–589.

